# Climbing Legumes: An Underutilized Resource With Significant Potential to Intensify Farming on Terrace Walls (FTW) for Smallholder Farmers

**DOI:** 10.1101/184952

**Authors:** Jaclyn C. Clark, Manish N. Raizada

**Affiliations:** Department of Plant Agriculture, University of Guelph, Guelph, ON Canada N1G 2W1

**Keywords:** legume, climbing, terrace agriculture, terrace riser, terrace wall, subsistence agriculture

## Abstract

Millions of subsistence farmers cultivate crops on terraces. These farmers face unique challenges including severe shortages of arable land and remoteness leading to poor access to inputs including nitrogen fertilizer. These challenges contribute to human and livestock malnutrition. Terrace walls (risers) as a vertical surface to grow climbing or trailing legumes represents an opportunity to help overcome these challenges. These crops are rich in minerals and protein, and their associated microbes produce nitrogen fertilizer. Rice bean is already grown on terrace risers in South Asia. This paper reviews the literature concerning crops that are currently farmed on terrace walls (FTW), then surveys climbing legume species that have potential for FTW, focusing on crops that are nutritious and tolerate shade (caused by the terrace wall) and resist drought (many terrace farms experience an extended dry season). A total of 29 legume species are discussed including climbing varieties of jack bean, common bean, cowpea, winged bean, horse gram and velvet bean. The review concludes by discussing the practical challenges of farmer adoption of FTW and makes concrete recommendations. Terrace wall cultivation of legumes represents an opportunity to intensify agriculture and increase resiliency in remote mountainous areas.

## 1. Introduction to terrace agriculture and its challenges

Subsistence smallholder agriculture generally refers to a farming system that uses few inputs on a very limited land base and produces food almost entirely for self-consumption (Graeub *et al*., 2016). A subset of subsistence farmers cultivate crops and raise livestock on steep hillside terraces. There appears to be no global estimates of the land area or number of farmers involved in terrace agriculture, an oversight that should be addressed by the Food and Agricultural Organization (FAO). Terraces have been reported to cover approximately 13,200,000 hectares in China and 2,000,000 hectares in Peru (Inbar and Llerena, 2000; Lu *et al*., 2009). Stanchi *et al*. noted that there is no reliable quantitative data concerning the total area currently covered by terraces in Europe (Stanchi *et al*., 2012). However it can be stated that terraced agriculture certainly covers a significant portion of land in Southeast Asia, the Himalayan Region, China, the Andes, Central America, East Africa, and a few locations in Europe. It is reasonable to estimate that minimally tens of millions of farmers worldwide rely on terraced land.

In general, subsistence farmers are inherently vulnerable to biophysical risks such as drought, flooding, pests and diseases (Morton, 2007). Such farmers may also face socioeconomic constraints, including but not limited to, restricted access to markets and political instability (Morton, 2007). Subsistence farmers who practice hillside terraced agriculture face additional unique challenges including severe limits on arable surface area, drudgery associated with walking up and down hillsides, the narrowness of fields which limit livestock labour, enhanced vulnerability to climate, erosion of soil from sloped topography, and reduced access to inputs and markets due to the inherent remoteness of such farms in mountainous regions, all of which combine to exacerbate human and livestock malnutrition (Chapagain and Raizada, 2017).

Terrace farmers must produce food on a very limited surface area. For example, in Nepal, the average agricultural landholding on the flat land (terai) is 1.26 ha, but that number shrinks to 0. 77 ha for hilly regions and 0.068 ha on mountainous land (Adhikary, 2004). Population pressure makes the task of producing enough yield to provide for a household with limited landholdings an increasingly difficult one (Paudel, 2002). For this reason, there is a need for terrace farmers to intensify production using the entire surface area available.

There is increased drudgery associated with terrace farming. Hillside farmers are constantly working against the rugged terrain and complex topography of their land. The narrowness of some terraces and steep terrain can limit access to livestock or machinery, resulting in increased human labour (e.g. land preparation) (Adhikary, 2004). Furthermore, farmers need to walk up and down steep grades with heavy loads, which places particular hardship on women, for not only is hill agriculture dependant on their labour, but they are also traditionally tasked with household duties and childcare (Pande, 1996).

The interplay between mountainous topography and climate exacerbates the vulnerability of subsistence farmers. The varying topography creates microclimates and diverse soil characteristics over small areas (Upadhyay, 1993), which complicates the development of best management practices and limits where and when a particular crop can be grown within a household’s already limited land holdings (Chapagain and Raizada, 2017; Pande, 1996). Many terrace farms are located in the sub-tropics which have uni-modal or bi-modal rainfall patterns resulting in an extended dry season which severely limits production (Small and Raizada, 2017). Terrace farmers are made more vulnerable by global climate change. For example, in Nepal, climate change has been associated with more frequent severe weather events (flooding, hailstorms, drought, delayed Monsoon rainfall) (Small and Raizada, 2017).Global climate change is expected to decrease the predictability of rainfall, and warming will also shift spring melting earlier (Morton, 2007). In mountainous regions where meltwater can be used for irrigation, this phenomenon will leave less available during the dry summer months when it is critically needed (Morton, 2007).

Another problem concerning terrace agriculture is soil stability. It was found in the mid-hills of Nepal that 93% of farmers faced some amount of terrace failure, which they spent an average of 14 days of labour per year repairing (Gerrard and Gardner, 2000). Preventing rain from directly hitting the terraces, along with root systems that stabilize soils at terrace edges, may help to prevent terrace failure (Acharya et al., 2007; Andersen, 2012; Gerrard and Gardner, 2000; Van Dijk and Bruijnzeel, 2004). Erosion from terrace edges, causing loss of nutrient-rich topsoil, is an important consequence of terrace topography (Van Dijk and Bruijnzeel, 2004). For example, in a study conducted over 2 years in the mid-hills of Nepal, there were losses of up to 12.9 tonnes of soil per hectare per year (Gardner and Gerrard, 2003). Conservation methods such as strip cropping have been attempted with variable success, however farmers are unwilling to sacrifice cultivatable land unless there will be a tangible benefit in terms of income (Acharya *et al*., 2008).

Remoteness can compound the impact of soil degradation and low productivity by making access to restorative agricultural inputs difficult. For example, on terrace farms in the mid and high hills of Nepal, limited income and remoteness has been shown to prevent local farmers from having access to markets to purchase inputs such as nitrogen fertilizer to help replenish lost nutrients, in comparison to the foothills and flatter terai region (Paudyal et al., 2001). Practices or crops that allow farmers to be self-sufficient in soil nutrient management are much needed. Reduced access to markets prevents farmers from gaining cash income from their products, and reduces access to knowledge, exemplified in Paudyal *et al*.’s review of maize in Nepal, which notes that sale of surplus grain and vegetables and access to extension are limited in the remote high hills (Paudyal *et al*., 2001). Vulnerability to emergencies is also increased as observed in the recent 2015 Nepal earthquake (Neupane, 2015). As mentioned above, terrace farmers in many regions experience an extended dry season; this causes seasonal out migration of men or entire families in search of work, which is made increasingly challenging in remote areas due to poor access to transportation and communication infrastructure, often along with issues of labour exploitation and discrimination as highlighted in a review of seasonal migration in Ethiopia (Asfaw *et al*., 2010). Any new approach that can generate on-farm income during the dry season may help to prevent this social upheaval.

All of the above factors contribute to human and livestock malnutrition amongst terrace farm households (Chapagain and Raizada, 2017). In general, smallholder farmers may have adequate calories, but lack some essential nutrients, of which protein (amino acid) deficiency and iron deficiency (anemia) are particularly problematic (Broughton et al., 2003; WHO, 2001). The latter causes weakness and lessens the ability of farmers to work (WHO, 2001). It is estimated that 50% of pregnant women and 40% of preschool aged children in developing nations are anemic (WHO, 2001). Zinc deficiency is also characteristic of people who tend to get most of their calories from cereals, and have minimal high-quality protein (Darton-Hill, 2013). Zinc deficiency can cause stunted growth, low immune system functioning and diarrheic disease (Darton-Hill, 2013). Other significant micronutrient deficiencies identified by the World Health Organization include vitamin A, folate, and iodine; these often occur because of inadequate diversity in the diet due to limited resources (Akhtar, 2016; Johns and Eyzaguirre, 2007).With respect to livestock, during the dry season it becomes difficult for farmers in many regions to find enough high quality fodder (Small and Raizada, 2017). Underweight livestock, overgrazing and illegal harvest of fodder from forests are just three of the several repercussions that arise from this situation (Upadhyay, 1993).

## 2. The potential for legume cultivation on terrace walls (risers)

Use of the vertical surface of terrace walls (risers, Fig. 1) to grow climbing or trailing legumes represents an opportunity to help overcome some of the challenges faced by terrace farmers. Legumes could be planted at either the base or top of the riser, and climb up or trail down the vertical surface, respectively. This technique may be referred to as farming on terrace walls (FTW). Growth of legumes on risers have the potential to help terrace farmers intensify their farms without the need for additional space, help improve their cash incomes, gain resiliency to drought, reduce soil erosion, improve soil fertility without external nitrogen fertilizer, and improve human and livestock nutrition.

**Figure 1.**
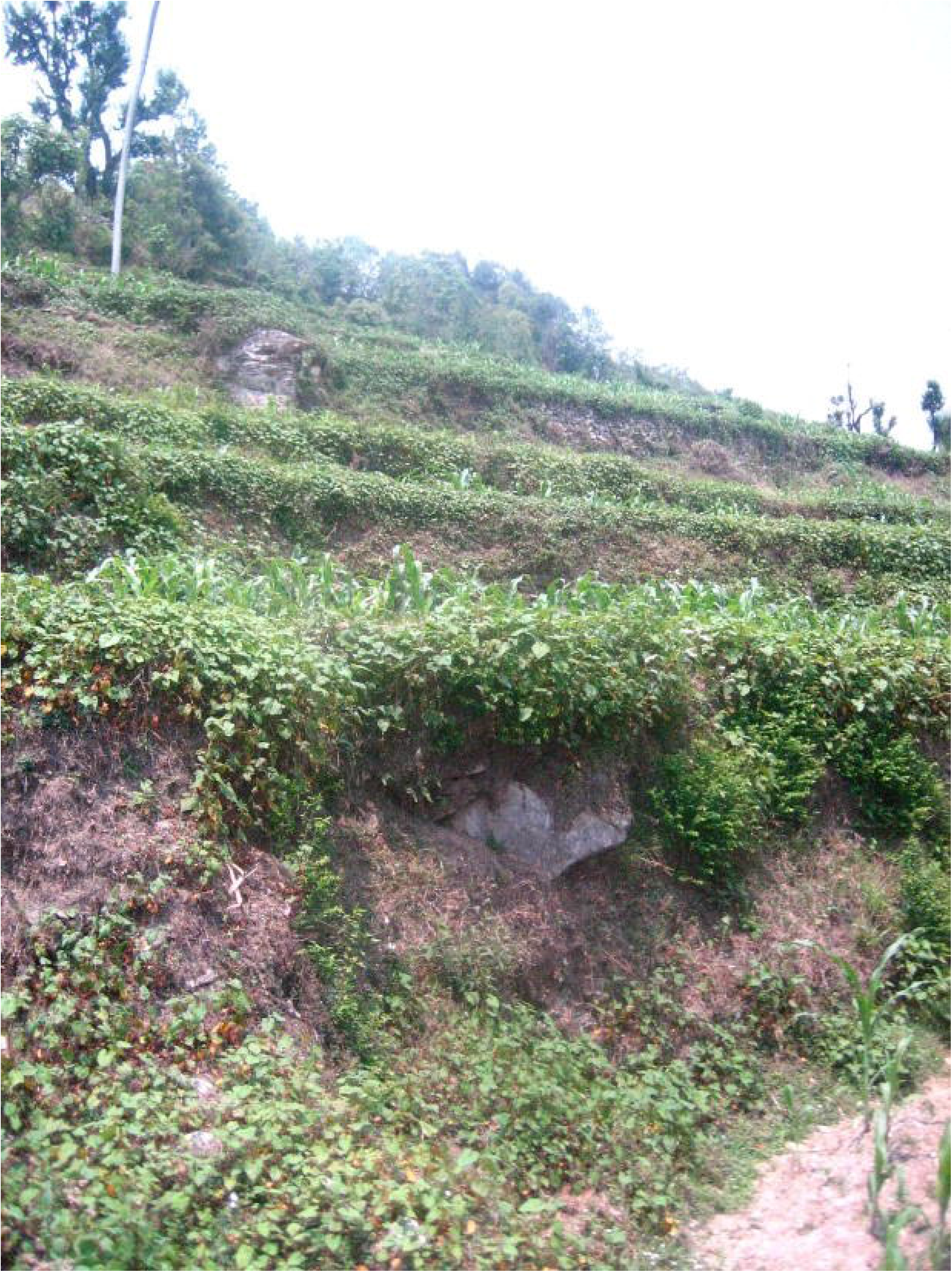
Example of terrace risers in Nepal as an underutilized surface area with potential to grow crops (photo credit: Manish Raizada).

Legumes belong to the family Leguminosae (or Fabaceae) which consists of over 750 genera and 20,000 species (Upadhyaya *et al*., 2011). They range from small herbs to large trees, however the relevant types for FTW are edible legumes with climbing and trailing characteristics. These legumes include several major edible human crops such as common beans, peas and cowpeas, but also underutilized and wild legumes described in this review. However, the legumes also include cover crops and green manures that, aside from soil benefits, can be used as livestock fodder. Certain legumes are able to climb because they have specialized structures called tendrils, a type of modified aerial stem (Tortora and Parish, 1970). Leaflets are replaced by tendrils which may vary in structure, but all function to support the stem of the plant (Langenheim, 1982). They are long and slender, allowing them to wind around objects they encounter, and some have small disks that stick on to supporting structures (Tortora and Parish, 1970). Legume grains generally contain high levels of protein, carbohydrates, fibre and minerals (iron and zinc) that make them an important tool for combating human malnutrition (Broughton *et al*., 2003; Upadhyaya *et al*., 2011). The nutritional characteristics of legume leaves also make them a protein-rich fodder for livestock (Upadhyaya *et al*., 2011). The issue of fodder shortage in the dry season may be resolved if drought tolerant legumes are selected that continue to produce fodder using only residual moisture (Small and Raizada, 2017). The high protein content of legumes is due to their unique ability to associate with symbiotic bacteria (rhizobia) that inhabit root nodules (Dilworth *et al*., 2008); these bacteria convert atmospheric nitrogen gas (N2) into ammonia (NH3) which serves as a limiting building block for amino acids, including those that lead to human protein malnutrition (Broughton *et al*., 2003). This process is called biological nitrogen fixation (Dilworth *et al*., 2008). Legume leaves and roots also have enhanced levels of plant-available nitrogen and protein, which, if not harvested, can be incorporated into soils as a form of organic nitrogen (Graham and Vance, 2003). Legume-derived nitrogen reduces the need for synthetic nitrogen fertilizer which would require cash and access to markets, which are both limited due to the remoteness of terrace farms, as noted above. Incorporation of legume residues into soil can increase soil organic matter and structure, to improve nutrient holding capacity and prevent erosion (Dilworth *et al*., 2008). Spreading-type legumes can also provide physical coverage to soils against rainfall, thus preventing erosion (Dilworth *et al*., 2008), especially if these varieties are planted at terrace edges.

## 3. Scope of this review

This paper reviews the limited literature concerning the current cultivation of crops and forages on terrace walls. After defining ideal agronomic traits, we survey climbing legume species that have potential to grow on terrace walls, focusing on crops that tolerate stress such as drought, are shade tolerant and are nutritious to humans and/or livestock. The review discusses the practical challenges that will limit farmer adoption of this practice. The paper concludes by making concrete recommendations to address these challenges.

## 4. Literature concerning current cultivation on terrace walls (terrace risers)

There has been no holistic assessment of terrace wall cultivation, and it is difficult to determine through the literature where the idea may have originated. One could infer from this that it was developed by indigenous farmers through trial and error. From the scattered reports that exist on terrace wall cultivation, Andersen briefly notes that short varieties of rice bean (Vigna umbellata) have been observed to grow on risers in India and Nepal to provide soil stability, feed and fodder (discussed in detail below) (Andersen, 2012). Most papers have noted in passing that grasses or forages are grown on risers as a strategy for increased structural stability or for improved nutrient health of terrace soils (Acharya *et al*., 2007; Andersen, 2012; Gerrard and Gardner, 2000; Pilbeam *et al*., 2000; Sharma *et al*., 2001). The lessons from these studies may help to inform efforts to cultivate legumes on terrace walls.

For example, Pilbeam *et al*. reported that fodder grasses can be grown on terrace risers in the mid-hills of Nepal; these riser grasses could contribute to the portion of livestock diets (30-45%, regardless of species) that is comprised of grass (Pilbeam *et al*., 2000). It was reported that when the grass was removed from risers and fed to livestock, and if the resulting livestock manure was spread on terrace benches, then there was a net movement of nitrogen from non-agricultural to agricultural land (Pilbeam *et al*., 2000). The fact that risers were designated as “non-agricultural” space indicates that the vertical area is generally not thought of as productive or useful in terms of cultivation.

Gerrard and Gardner noted in their study of landslide events in the mid-hills of Nepal that terraces were more susceptible to failure early in the season when irrigation was applied and riser vegetation was not yet established (Gerrard and Gardner, 2000). The lesson learned from this study was that if structural stability is one of the goals of planting on risers, fast growing varieties would most likely be more effective. In India, the National Watershed Development Project for Rain-fed Areas distributed broom grass (Thysanolena maxima) to farmers, hoping to utilize its soil binding properties for terrace stability (Sharma *et al*., 2001). In this case study, Sharma *et al*. found that perennial species were more helpful in preventing structural breakdown of terrace risers than annuals (Sharma *et al*., 2001).

A study by Acharya *et al*. may be especially informative in the context of farmer adoption of a new crop variety or practice. In order to reduce nutrient losses in the mid-hills of Nepal, the researchers planted fodder grass (Setaria anceps) on risers (Acharya *et al*., 2007). It was concluded that this practice prevented runoff but had no effect on the more significant problem of leaching (Acharya *et al*., 2007). The relevant observation from this study was that farmer adoption was more likely out of interest for higher quality fodder than environmental improvements, and that the production of fodder on-farm saved time over collecting it from the forest, thus reducing labour (Acharya *et al*., 2007). It was noted that establishment of the grass on risers initially caused more disturbance to the farming system, with the major benefits observed in subsequent years, which may be an obstacle to adoption (Acharya *et al*., 2007). This study emphasized the importance of farmers’ priorities as vital starting points for introducing any new species (Acharya *et al*., 2007). The study also discusses the importance of multiple decision-making factors when it comes to the promotion of farmer adoption of a new technique (Acharya *et al*., 2007).

## 5. Climbing legume agronomic traits that address the vulnerabilities of terrace farming systems

Terrace farming systems are diverse around the world, in terms of their biophysical and socioeconomic contexts, and hence a particular plot may have unique priorities in terms of crop species selection. However, amongst the climbing legumes, there are some species that may be able to address the above noted vulnerabilities of many terrace farming systems based on their agronomic traits. These traits are summarized (Fig. 2). Learning from earlier studies, to ensure adoption, a climbing legume must have obvious utility as food, fodder, or for income generation. Secondary traits would include: drought tolerance (to provide nutritious food and fodder in the dry season), shade tolerance (since the terrace wall may cause shading), adaptation to low chemical inputs (to combat remoteness), low labour requirements (to reduce drudgery), and utility as a cover crop or green manure (to maintain and improve soil quality). Tertiary traits include those that allow crop growth under marginal soils (saline, acidic, alkaline) or for which improved varieties have been bred.

**Figure 2.**
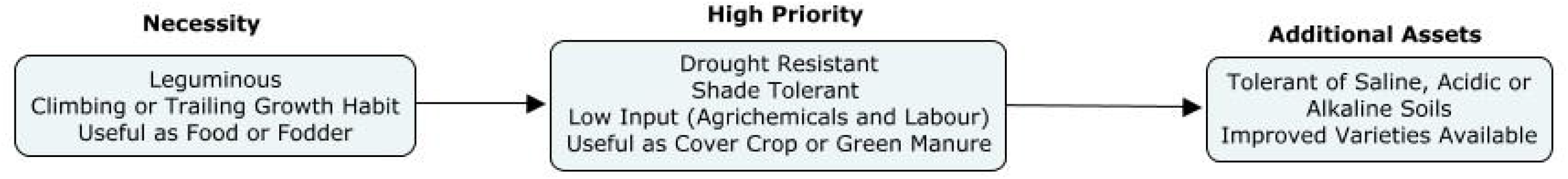
Agronomic traits of crops that address the vulnerabilities of terrace farming systems

## 6. Survey of climbing legume species that have potential to mitigate the vulnerabilities of terrace farming systems

The current literature was reviewed to identify species with agronomic, nutritional and alternative use traits that may render them successful for growth on terrace walls. In total, 29 climbing legume species belonging to 16 different genera show some promise with respect to farming on terrace walls (complete list, Table S1). The species are organized by their reported tolerance to abiotic stress (Table 1), rainfall requirements for those that are drought tolerant (Table 2), reported utility (Table 3), specific nutritional details (Table 4), and current level of genetic improvement (Table 5). Below is a summary of each candidate genus:

**Table 1.** Climbing legumes with respect to reported tolerance to abiotic stress

| Genus | Species | Common name | Reported Tolerance to Unfavourable Conditions |  |  |  |  |
| --- | --- | --- | --- | --- | --- | --- | --- |
|  |  |  | Drought | Shade | Poor Soil | pH | Flooding |
| *Amphicarpaea* | *A. bracteata* | Hog Peanut |  | ● |  |  |  |
|  | *A. edgeworthii* | N/A |  |  | ● (calcareous) |  |  |
| *Apios* | *A. americana* | Groundnut |  |  |  | ● (low) | ● |
|  | *A. priceana* | N/A |  |  |  | ● (low and high) |  |
| *Canavalia* | *C. ensiformis* | Jack Bean | ● (once established) | ● | ● (saline) |  | ● |
|  | *C. gladiata* | Sword Bean | ●* (once established) |  |  |  |  |
| *Cyamopsis* | *C. tetragonoloba* | Cluster Bean | ● |  | ● (saline) | ● (high) |  |
| *Lablab* | *L. purpureus* | Hyacinth Bean | ● (once established) | ● |  |  |  |
| *Lathyrus* | *L. sativus* | Grass Pea | ● |  | ● (low fertility) | ● (high) | ● |
| *Macrotyloma* | *M. uniflorum* | Horse Gram | ● |  |  |  |  |
| *Mucuna* | *M. pruriens* | Velvet Bean | ● * |  |  | ● (low) |  |
| *Pachyrhizus* | *P. tuberosus* | Ashipa |  |  |  | ● (low) |  |
| *Phaseolus* | *P. lunatus* | Lima Bean | ● |  |  |  |  |
|  | *P. vulgaris* | Common Bean | ● * |  |  |  |  |
| *Pisum* | *P. sativum* | Field/Green Pea | ● * |  |  |  |  |
| *Psophocarpus* | *P. tetragonolobus* | Winged Bean |  |  | ● (low OM) |  |  |
| *Pueraria* | *P. lobata* | Kudzu | ● |  |  |  |  |
|  | *P. phaseoloides* | Tropical Kudzu | ● | ● (moderate) |  |  |  |
|  | *P. tuberosa* | Indian Kudzu |  |  | ● (eroded or exposed) |  |  |
| *Sphenostylis* | *S. stenocarpa* | African Yam Bean |  |  | ● (low fertility) | ● (low) |  |
| *Vigna* | *V. umbellata* | Rice Bean | ● |  |  |  |  |
|  | *V. unguiculata* | Cowpea | ●* | ● | ● (salinity) | ● (low) |  |

\*tolerant varieties available

^*^tolerant varieties available

**Table 2.** Rainfall and distribution requirements of drought tolerant climbing legumes.

| **Genus** | **Species** | **Common Name** | **Reported Specific Water Requirement/**<br>**Drought Capability** |
| --- | --- | --- | --- |
| *Canavalia* | *C. ensiformis* | Jack Bean | Requires well-distributed 900-1200 mm/year to establish, after that can be successful as low as 650 mm/year |
|  | *C. gladiata* | Sword bean | Requires well-distributed 900-1500 mm/year initially, some varieties drought resistant once established |
| *Cyamopsis* | *C. tetragonoloba* | Cluster Bean | Grows well with annual rainfall of 500-750 mm/year |
| *Lablab* | *L. purpureus* | Hyacinth Bean | Distribution dependant, can grow with 600-900mm/year. Requires adequate rainfall for first 2-3 months after planting, then becomes drought resistant and can produce well into the dry season |
| *Lathyrus* | *L. sativus* | Grass Pea | Prefers rainfall of 350-600 mm/year |
| *Macrotyloma* | *M. uniflorum* | Horse Gram | Rainfall requirement can be as low as 200 mm/year |
| *Mucuna* | *M. pruriens* | Velvet Bean | Flourishes at 1200-1500 mm/year, however there are some drought resistant varieties available |
| *Phaseolus* | *P. lunatus* | Lima Bean | Reasonably drought tolerant, however also successful with high rainfall. Can grow with anywhere between 500-1500+ mm/year. |
|  | *P. vulgaris* | Common Bean | Significant attention in breeding programs. Some areas seen success with 300-400 mm/year |
| *Pisum* | *P. sativum* | Field/Green Pea | Preferably 800-1000 mm/year, however some success with as low as 400 mm/year |
| *Pueraria* | *P. lobata* | Kudzu | Thrives with more than 1000 mm/year, however also drought resistant |
|  | *P. phaseoloides* | Tropical Kudzu | Reasonably tolerant of drought |
| *Vigna* | *V. umbellata* | Rice Bean | Optimum yields between 1000-1500 mm/year, however also tolerant of drought |
|  | *V. unguiculata* | Cowpea | Rainfall requirement depends on duration type; quick maturing varieties can grow with less than 600 mm/year |

**Table 3.** Uses of candidate climbing legume species

| Genus | Species | Common Name | Reported Utility |  |  |  |  |
| --- | --- | --- | --- | --- | --- | --- | --- |
|  |  |  | Food | Forage or Fodder* | Green manure | Cover crop | Feed* |
| *Amphicarpaea* | *A. bracteata* | Hog Peanut | ● | ● |  |  |  |
| *Apios* | *A. americana* | Groundnut | ● |  |  |  |  |
|  | *A. priceana* | N/A | ● |  |  |  |  |
| *Canavalia* | *C. ensiformis* | Jack Bean | ● | ● | ● | ● |  |
|  | *C. gladiata* | Sword Bean | ● | ● | ● | ● |  |
| *Cyamopsis* | *C. tetragonoloba* | Cluster Bean | ● | ● | ● |  |  |
| *Lablab* | *L. purpureus* | Hyacinth Bean | ● | ● | ● |  |  |
| *Lathyrus* | *L. sativus* | Grass Pea | ● |  |  |  | ● |
|  | *L. tuberosus* | Earthnut Pea | ● | ● |  |  |  |
| *Macrotyloma* | *M. uniflorum* | Horse Gram | ● | ● | ● |  |  |
| *Mucuna* | *M. pruriens* | Velvet Bean | ● |  | ● | ● | ● |
| *Pachyrhizus* | *P. tuberosus* | Ashipa | ● |  |  |  |  |
| *Phaseolus* | *P. coccineus* | Runner Bean | ● |  |  |  |  |
|  | *P. lunatus* | Lima Bean | ● |  | ● |  | ● |
|  | *P. vulgaris* | Common Bean | ● |  |  |  | ● |
| *Pisum* | *P. sativum* | Green/Field Pea | ● | ● |  |  | ● |
| *Psophocarpus* | *P. tetragonolobus* | Winged Bean | ● | ● |  | ● |  |
| *Pueraria* | *P. lobata* | Kudzu | ● | ● | ● |  |  |
|  | *P. phaseoloides* | Tropical Kudzu | ● | ● | ● | ● |  |
|  | *P. tuberosa* | Indian Kudzu | ● |  |  |  | ● |
|  | *P. americana* | American Kudzu | ● |  |  |  |  |
| *Sphenostylis* | *S. stenocarpa* | African Yam Bean | ● |  |  |  |  |
| *Vicia* | *V. sativa* | Common Vetch | ● | ● |  |  |  |
| *Vigna* | *V. umbellata* | Rice Bean | ● | ● | ● |  |  |
|  | *V. unguiculata* | Cowpea | ● |  | ● | ● |  |

\*forage and fodder refer to unprocessed animal feed (e.g. pasture grazing, cut and carry) whereas feed refers to processed or refined animal feed

^*^forage and fodder refer to unprocess al feed (e.g. pasture grazing, cut and carry) whereas feed refers to processed or refined animal feed

**Table 4.** Edibility and nutritional information of climbing legumes

| Genus | Species | Common Name | Additional nutritional notes |
| --- | --- | --- | --- |
| *Amphicarpaea* | *A. bracteata* | Hog Peanut | Sub-terrain seeds eaten roasted. |
| *Apios* | *A. americana* | Groundnut | Tubers |
|  | *A. priceana* | N/A | Slightly larger tubers than *A. americana*, poor amino acid profile. |
| *Canavalia* | *C. ensiformis* | Jack Bean | Mature seeds eaten, must be boiled first to soften and get rid of toxic components. May be roasted as a coffee substitute. Immature pods boiled and eaten as a vegetable. |
|  | *C. gladiata* | Sword Bean | See *C. ensiformis*. |
| *Cyamopsis* | *C. tetragonoloba* | Cluster Bean | Immature pods boiled, fried, or processed into gum. Leaves sometimes cooked as a vegetable. |
| *Lablab* | *L. purpureus* | Hyacinth Bean | Pods used as vegetable or in curries. Beans may be boiled or roasted, sometimes fried into a cake. |
| *Lathyrus* | *L. sativus* | Grass Pea | Beans consumed, however overconsumption associated with degeneration of upper motor neurons due to neurotoxin. Advised to be eaten in moderation and paired with antioxidant rich foods. |
|  | *L. tuberosus* | Earthnut pea | Tubers eaten raw or roasted. Seeds sometimes consumed. |
| *Macrotyloma* | *M. uniflorum* | Horse Gram | Seeds boiled, fried or made into cakes. |
| *Mucuna* | *M. pruriens* | Velvet Bean | Beans must be soaked or boiled to remove toxic component. Sometimes eaten roasted. |
| *Pachyrhizus* | *T. tuberosus* | Ashipa | Roots eaten raw or cooked, tubers may be used to make custard, pudding or flour. Young pods cooked, must be boiled to get rid of insecticidal toxin. |
| *Phaseolus* | *P. coccineus* | Runner Bean | Immature pod and seeds within boiled. Mature seeds eaten fresh or dried, tubers sometimes used for starch. |
|  | *P. lunatus* | Lima Bean | Mature dry beans can be boiled, fried, baked or ground into flour. Immature pods can be eaten as a vegetable. |
|  | *P. vulgaris* | Common Bean | Considerable variation in beans, often mixed with carbohydrate source (e.g. rice/cassava). |
| *Pisum* | *P. sativum* | Field/Green Pea | Wide variation. Seeds eaten dried or fresh. Some immature seedpods consumed. Many processing options, usually canning. |
| *Psophocarpus* | *P. tetragonolobus* | Winged Bean | Full plant edible: young pods (raw, boiled, steamed, fried or pickled), leaves, shoots (similar to asparagus), flowers (steamed or fried), seeds, and tubers. |
| *Pueraria* | *P. lobata* | Kudzu | Leaves, shoots and flowers consumed. Roots harvested for starch. |
|  | *P. phaseoloides* | Tropical Kudzu | Tuberous roots edible, however not significantly recorded. |
|  | *P. tuberosa* | Indian Kudzu | Tubers considered a famine food. |
| *Sphenostylis* | *S. stenocarpa* | African Yam Bean | Seeds can be boiled, soaked and fried, tubers usually boiled or roasted. |
| *Vicia* | *V. americana* | American Vetch | Young shoots, immature seedpods and mature seeds consumed (traditionally by Native Americans). |
|  | *V. sativa* | Common Vetch | Beans may be consumed by humans, not very digestible. |
| *Vigna* | *V. umbellata* | Rice Bean | Beans usually mixed with rice. Pods and green seeds cooked as a vegetable. |
|  | *V. unguiculata* | Cowpea | Beans in soups and cakes, immature seeds and pods may be eaten as a vegetable. Shoots can be boiled. |

**Table 5.** Current status of genetic development of climbing legumes for agricultural use.

| Well Established as Crop --<br>Potential for Genetic<br>Improvement |  | Limited Cultivation – Will<br>Benefit Substantially from<br>Further Breeding |  | Only Wild Growth/Gathering –<br>Requires Domestication |  |
| --- | --- | --- | --- | --- | --- |
| Species | Common<br>Name | Species | Common<br>Name | Species | Common<br>Name |
| *Lablab purpureus* | Hyacinth Bean | *Amphicarpaea bracteata* | Hog Peanut | *Amphicarpaea africana* | N/A |
| *Macrotyloma uniflorum* | Horse Gram | *Apios americana* | Groundnut | *Amphicarpaea edgeworthii* | N/A |
| *Phaseolus coccineus* | Runner Bean | *Apios priceana* | N/A | *Lathyrus japonicas* | Sea Pea |
| *Phaseolus lunatus* | Lima Bean | *Canavalia ensiformis* | Jack Bean | *Lathyrus tuberosus* | Earthnut Pea |
| *Phaseolus vulgaris* | Common Bean | *Canavalia gladiata* | Sword Bean | *Pueraria phaseoloides* | Tropical Kudzu |
| *Pisum sativum* | Green Pea | *Cyamopsis tetragonoloba* | Cluster Bean | *Pueraria americana* | Indian Kudzu |
| *Vigna umbellata* | Rice Bean | *Lathyrus sativus* | Grass Pea | *Vicia sativa* | Common Vetch |
| *Vigna unguiculata* | Cowpea | *Mucuna pruriens* | Velvet Bean |  |  |
|  |  | *Pachyrhizus tuberosus* | Ashipa |  |  |
|  |  | *Psophocarpus tetragonolobus* | Winged Bean |  |  |
|  |  | *Pueraria lobata* | Kudzu |  |  |
|  |  | *Sphenostylis stenocarpa* | African Yam Bean |  |  |

### Amphicarpaea

There are three noteworthy species of climbing legumes from the genus Amphicarpaea, a close relative of the soybean family. These species generally grow on separate continents: A. africana, A. bracteata (also referred to as hog peanut or talet bean) and A. edgeworthii from Africa, North America and Asia, respectively, with A. edgeworthii adapted to the mid-altitude Himalayas (Turner and Fearing, 1964) where there are many terrace farms. They all have twining vines, however most available information focuses on the two most similar species, A. bracteata and A. edgeworthii (Turner and Fearing, 1964). Both of these species produce two types of fruit, aerial pods and subterranean beans, from heteromorphic flowers – a botanical feature that gave rise to the name of this genus (Graham, 1941; Zhang *et al*., 2006). The underground pods are primarily the ones consumed, and they have also been reported to be dug up and eaten by pigs, giving rise to the common name ‘hog peanut’ (Graham, 1941). A. bracteata interestingly showed >3-fold increased productivity under 80% shaded conditions compared to full sunlight (Lin *et al*., 1999), suggesting that it may perform well in the shadow of terrace walls. Though some of the characteristics of this genus sound promising, limited recent literature suggests that the species may need some genetic improvement.

### Apios

A. americana and A. priceana are vines native to North America, and have been similarly neglected in recent studies. Some older reviews cited their potential in agriculture as a crop and to enable soil improvement (Graham, 1941; Putnam *et al*., 1991). Both species have edible tubers that appear to have high levels of carbohydrates and protein, although it was found that the amino acid profile of A. priceana is relatively poor compared to major root crops (Walter *et al*., 1986). A. americana was able to produce nodules when inoculated with rhizobia traditionally used for soybean and cowpea (Putnam *et al*., 1991), indicating good potential for nitrogen fixation if introduced into agricultural systems.

### Canavalia

Canavalia ensiformis is a legume commonly referred to as jack bean or overlook bean, which has several agronomic characteristics and diverse uses that may render it useful to terrace farmers (Bazill, 1987; Haq, 2011; Kay, 1979). For this reason, this species will be reviewed extensively here. The crop is a weak perennial (because of this it is usually grown as an annual) native to the West Indies and Central America (Haq, 2011; Kay, 1979). C. ensiformis is adapted to the humid tropics, and though it is reported to require 900-1200 mm rainfall/year during early growth, once established it has a deep root system and becomes drought tolerant and can survive on as little as 650 mm/year (Haq, 2011; Kay, 1979; Pound *et al*., 1972). It has also been reported to be tolerant of saline or waterlogged soils (Haq, 2011; Kay, 1979), the latter condition observed on rice terraces as already noted. A study conducted in Costa Rica found that C. ensiformis produced well in shaded conditions, able to grow with only 18% of full sunlight (Bazill, 1987), making it a strong candidate for growth on terrace walls. Indeed, the shade tolerance of this species has allowed its widespread use as a cover crop under cocoa, coconut, citrus and pineapple (Haq, 2011).

Multiple authors have noted that agronomic data concerning this species is limited. It is planted both in rows and broadcast, with seeding rates and spacing data reported (Haq, 2011; Kay, 1979). C. ensiformis is often observed to be intercropped with other crops like sugarcane, coffee, tobacco, rubber and maize (Arim *et al*., 2006; Haq, 2011). In one study, maize that had been intercropped with C. ensiformis performed better when infested with Pratylenchus zeae, a pathogenic nematode (Arim *et al*., 2006). It was hypothesized by the author that either the toxic components in C. ensiformis created low phosphorus conditions which are unfavourable to P. zeae, or intercropping with the legume improved the growth of maize and hence made it more resistant to pests (Arim *et al*., 2006). In general, C. ensiformis has been reported to be pest resistant, perhaps because it produces hydrogen cyanide (HCN) and other toxic properties (Haq, 2011; Pound *et al*., 1972).

Pods are reportedly ready to harvest after 3-4 months, and the seeds after 6-10 months (Haq, 2011). Forage yields have been reported to be ~6000 kg/ha, and dry bean yields range from 1200-4800 kg/ha (Haq, 2011; Kay, 1979). A study conducted in the Dominican Republic noted that multiple cuts of forage can be removed in one season, and reported seed yield that overlapped with soybeans under the conditions grown (Pound *et al*., 1972).

There are many different reported utilities of C. ensiformis. The crop has good seed and storage qualities (Pound *et al*., 1972). Young leaves, pods and immature seeds are all edible by humans (Haq, 2011), however the seeds must be soaked or boiled for several hours to soften and remove toxic components, but even following these treatments, they are purportedly not very palatable (Kay, 1979). The plants can be a valuable forage for livestock, and sometimes dried seeds are used as feed, however poisoning has been reported if seeds are uncooked or comprise more than 30% of the livestock diet (Kay, 1979). Two reports also mentioned that the high protein content of C. ensiformis (generally reported between 23 and 28%) lends itself to the opportunity for processing into protein isolates (Haq, 2011; Kay, 1979). C. ensiformis has also been studied as a green manure, and it was found that the deep root system and exceptional nodulation gave the species a high capacity to provide nitrogen to subsequent crops (Wortmann *et al*., 2000). The species was found to fix 133 kg N/ha from the atmosphere, and was generally more effective than several other legumes, including soybeans (Wortmann *et al*., 2000).

There is limited information concerning genetic resources and improvement of C. ensiformis, though there are apparently some breeding programs in India, Indonesia and Africa with goals of creating higher yielding and lower toxicity varieties (Haq, 2011).

In the same genus, Canavalia gladiata, a more vigorous perennial climber often called the sword bean, has some similarly encouraging characteristics. Originally cultivated in India (Rajaram and Janardhanan, 1992) it was reportedly spread around the world by ancient peoples intrigued by the sword-like length of its seed pod (Herklots, 1972). Though it requires high temperatures and reasonably fertile soils, there are some varieties that are resistant to drought once established, and reports suggest that it is fairly resistant to pests and diseases (Ekanayake *et al*., 2000). Yields of sword bean overlap with the range of yields observed with soybeans, and this species has diverse uses as summarized (Table 3) (Ekanayake *et al*., 2000). Since it is already currently cultivated throughout Asia, acquiring seed and establishing growing methods should be easier than for some of the lesser known species discussed in this article.

### Cyamopsis

As discussed earlier in this review, legumes are versatile plants with a multitude of uses. A good example of that is Cyamopsis tetragonoloba, commonly known as the cluster bean or guar in India and Pakistan where it is already a major food crop. Kumar *et al*. (2005) have extensively reviewed the Indian literature concerning this crop, including selection and breeding efforts (Kumar, 2005). The beans of this plant, a drought tolerant annual, may be eaten as a vegetable, but it also produces a valuable gum (Kay, 1979; Whistler and Hymowitz, 1979). Once processed, guar gum can be used for applications both in the food industry and in other types of manufacturing (Mudgil *et al*., 2014). It is also reported that the saline and alkaline tolerating properties of this species make it useful in reclamation of degraded soil (Kay, 1979).

### Lablab

One of the most versatile climbing legumes included in this review is Lablab purpureus, a climbing perennial that has over 150 common names (Maass *et al*., 2010). It appears to have originated in Africa and today continues to be grown in the highlands of East Africa (Haque and Lupwayi, 2000) with reports that it was grown in India as early as 1500 BC (Maass and Usongo, 2007), though today it is grown worldwide (Maass *et al*., 2010). It is considered to be an excellent green manure to support the growth of cereals in a rotation system (Haque and Lupwayi, 2000; Wortmann *et al*., 2000) though farmers appear to adopt it primarily when used as a livestock forage in the dry season (Haque and Lupwayi, 2000). This species shows great diversity in both agro-morphological characteristics and potential uses, with possibly 3000 accessions available for future crop improvement (Maass *et al*., 2010). It tends to require adequate water during the early stages of growth, but once established can be extremely drought resistant and produce many edible parts including pods, beans and leaves (Haq, 2011; Maass *et al*., 2010). One of the most promising qualities of this crop is that it has undergone some genetic development to create improved varieties, significantly in India and Bangladesh, though yields are generally considered to be low (Maass *et al*., 2010).

### Lathyrus

Another crop that shows potential for harsh conditions is the species Lathyrus sativus, an annual, straggling crop present throughout most of Asia and some parts of Africa, including Ethiopia (Hillocks and Maruthi, 2012). This species is commonly referred to as grass pea. In South Asia, it is grown as a relay crop following rice (Hillocks and Maruthi, 2012). It has been reported that grass pea may be the oldest domesticated crop in Europe, originating from Spain (Hanbury *et al*., 2000; Pena-Chocarro and Pena, 1999). The crop is reported to tolerate a wide range of difficult conditions, and is considered a safety net for farmers during drought and flooding when other crops fail (Hillocks and Maruthi, 2012; Malek *et al*., 2000); waterlogging is common to rice terraces containing clay soils. However, the edible parts of this crop also contain a neurotoxin that, when consumed in large quantities, can cause a condition called ‘lathyrism’ in both humans and animals (Hanbury *et al*., 2000; Hillocks and Maruthi, 2012). The toxin may cause symptoms such as weakness and paralysis, thus reducing farm labour capacity, and this toxin is associated with several species of this genus (Hanbury *et al*., 2000). Other species that show potential include Lathyrus japonicas, commonly referred to as sea pea, and Lathyrus tuberosus, often referred to as the earthnut pea. Little is known about these two species, but there are claims that they were used historically for food and may have potential for similar uses to L. sativus (PFAF, 2012). The neurotoxin related to this genus and associated neurodegenerative condition is certainly an obstacle that makes adoption of these species unattractive to growers, with its seed being banned in some nations; however if improved varieties with low toxin levels can be developed from the >4000 accessions available for breeding (Hillocks and Maruthi, 2012), then the grass, sea and earthnut pea may be beneficial choices for challenging farming environments.

### Macrotyloma

Macrotyloma uniflorum, a multi-use legume known as horse gram, grows in parts of Asia and Africa, particularly in India where it was likely domesticated and has been found since ancient times (Mehra and Upadhyaya, 2013). This species is also sometimes referred to as Dolichos uniflorus, and is utilized as a low-grade pulse crop, a forage for cattle/horses (particularly because it is available throughout the dry season) and as a green manure (Mehra and Upadhyaya, 2013; Siddhuraju and Manian, 2007; Cook *et al*., 2005). It is widely cultivated, however limited attention has been paid to it in terms of genetic development or marketing, so it is still referred to as underutilized, similar to several other crops in this review. Nevertheless, horse gram shows promise because of its drought tolerance and nutritional qualities, along with the fact that it requires few inputs (Bravo *et al*., 1999; Mehra and Upadhyaya, 2013; Siddhuraju and Manian, 2007; Witcombe *et al*., 2008). It has been shown to be a good source of protein and carbohydrates, and potentially also iron and calcium as long as certain preparation methods are used to break down anti-nutritional compounds (Bravo *et al*., 1999; Sudha *et al*., 1995). Participatory trials have been conducted that showed considerable success at addressing some of the challenges of resource poor farmers when horse gram was intercropped with maize in India (Witcombe *et al*., 2008). Farmers reported that labour demand decreased due to ground cover provided (particularly female drudgery such as weeding), and they were able to harvest both grain for food and fodder for livestock from the crop (Witcombe *et al*., 2008). There have also been studies conducted that showed improvement when horse gram was intercropped with finger millet (Pradhan, *et al*., 2014). One study was based in the hilly regions of India which found increased yields and improved nitrogen and phosphorus status with the addition of horse gram (Narendra *et al*., 2010). There have been some genetic improvements with a focus on increasing yield and disease resistance (Bhardwaj *et al*., 2013), however further efforts appear to be required to make more seed available to farmers and to develop a market for the grain. Recently this species has received increased attention and been reviewed more extensively for its potential as a health food and nutraceutical (Prasad and Singh, 2015).

### Mucuna

The genus Mucuna contains one herbaceous climbing vine of potential interest to terrace farmers, M. pruriens, commonly referred to as velvet bean (Haq, 2011; Kay, 1979). This species, which originated in Asia, can be either an annual or perennial, and is now grown throughout the tropics, particularly in the western hemisphere (Haq, 2011; Kay, 1979). It is generally suited to high rainfall areas, however some drought tolerant varieties are reportedly available (Kay, 1979).

In southeast Asia, the immature pods and leaves are reportedly consumed, whereas in parts of Asia and Africa, the seeds are typically roasted, fermented or used as thickeners in soups (Haq, 2011). M. pruriens shows promise in providing some essential components to the diets of the rural poor; the grain has crude protein levels of between 15.1 and 31.4%, as well as significant portions of unsaturated fatty acids, fibre and energy (Haq, 2011; Siddhuraju *et al*., 1996). Velvet bean also has potential to replace some of the soybean present in animal feed, which generally provides the majority of the protein content but must be imported to tropical regions (Chikagwa-Malunga *et al*., 2009). However, adoption of velvet bean cultivation has been somewhat limited due to the presence of HCN and other anti-nutritional factors which may decrease its digestibility (Rich and Teixeira, 2005; Siddhuraju *et al*., 1996). It has been reported, however, that proper processing and cooking methods involving heat can significantly decrease the levels of undesirable compounds (Haq, 2011; Siddhuraju *et al*., 1996). When investigating the harvest window in which nutrition for animal feed was optimized, it was found that between 110-123 days after planting, crude protein and fibre content remains constant, although dry matter continues to increase, and generally varies widely with different environmental factors such as rainfall (Chikagwa-Malunga *et al*., 2009).

M. pruriens has also been used as a cover crop and green manure, with some success. One study noted that when intercropped with corn there were decreased negative impacts by nematodes, similar to the impact of C. ensiformis discussed earlier (Arim *et al*., 2006). Timing of planting for use as a cover crop (particularly with maize) has been a subject of investigation, for the success of such a system depends on many factors (Lawson *et al*., 2007; Ortiz-Ceballos *et al*., 2015). Velvet bean planted soon after the sowing of maize sometimes had issues of lowering maize yield through competition for resources, with M. pruriens reportedly climbing maize stalks and shading the crop (Lawson *et al*., 2007). It will be interesting to test how these two crops perform when velvet bean is allowed to grow on the terrace wall, with maize cropped along the remainder of the terrace. When velvet bean was planted as a cover crop 6 weeks after maize was planted, it produced less ground cover, however the maize yields were higher and there was still significant weed suppression (Lawson *et al*., 2007). Recently, to investigate the issue of smothering, M. pruriens was used in rotation with maize as a relay crop in fallow seasons, instead of being intercropped, and it was an effective green manure, improving the fertility and structure of the soil (Ortiz-Ceballos *et al*., 2015).

It is encouraging to note that recent studies are being undertaken to address obstacles to the adoption of this legume, therefore potentially leading to a more thorough understanding of its agronomic characteristics (e.g. drought tolerance, cover crop and nutritive potential) and how they may be useful in addressing the challenges of terrace farmers.

### Pachyrhizus

The genus Pachyrhizus (known as yam bean, and in South America commonly as ashipa/ahipa) consists of four main crop species with climbing varieties, of which P. tuberosus and P. erosus are the most significant (Pena-Chocarro and Pena, 1999). P. tuberosus is a tropical perennial herbaceous vine that is cultivated on trellises; it was likely domesticated in the Peruvian Andes, although its origin has been difficult to map due to its extremely long reported history of continuous cultivation and consumption in South America (Sorensen *et al*., 1997; PFAF, 2012). The hallmark of this legume is that it produces a nutritious tuber(s) which is used as a substitute for cassava; but unlike cassava which has a toxic tuber, this tuber is usually eaten raw (Sorensen *et al*., 1997). P. tuberosus requires at least 16 weeks of growth to flower, however in soils with limited fertility it requires a longer period (8-15 months) to produce tubers (Sorensen *et al*., 1997). It is reportedly able to grow when planted at the end of the rainy season using only residual moisture (Sorensen *et al*., 1997). The roots and pods have both been described as edible, however they must be thoroughly cooked to remove rotenone, an insecticide (Sorensen *et al*., 1997). There is limited information concerning nutritional composition or agronomic practices, most likely due to the fact that it is traditionally a part of shifting cultivation systems and consumed within the community itself (Sorensen *et al*., 1997). P. erosus, however, has been cultivated on a larger scale for export and is grown in tropical regions of most continents (Sorensen *et al*., 1997). This species is similar to P. tuberosus in structure and is also grown for its tuberous roots (Sorensen *et al*., 1997). The larger-scale production of this species has allowed for the development some processing industries (Melo *et al*., 2003). Regarding these crops, Sorensen *et al*. have extensively reviewed older literature from the 1920’s-1940’s (Sorensen *et al*., 1997), while Ramos-de-la-Pena *et al*. have reviewed the scarce data available from more recent years (Ramos-de-la-Pena *et al*., 2013).

### Phaseolus

Perhaps the most well-known legume genus is Phaseolus, which includes species that have climbing varieties, such as P. coccineus (runner bean), P. lunatus (lima bean) and P. vulgaris (common bean). P. vulgaris genetically diverged into two populations and was simultaneously domesticated in the Andean and Mesoamerican regions 8000 years ago (Gaut *et al*., 2014). This species is considered one of the most important legume of the world’s poor, cultivated and consumed worldwide (Broughton *et al*., 2003; Kay, 1979). It provides as much as 1/3 of dietary protein in some regions of the world (Gaut *et al*., 2014). In pre-Columbian America, P. vulgaris was intercropped with maize which provided support for climbing, as part of the “Three Sisters” intercrop (Zhang *et al*., 2014). Similar to P. vulgaris, P. lunatus was domesticated in both the Andean and Mesoamerican regions in parallel (Motta-Aldana *et al*., 2010), while P. coccineus is thought to have originated solely in Mexico (Kay, 1979). P. lunatus is drought tolerant, producing beans with only 500-600 mm of rainfall, however P. coccineus is extremely drought susceptible [35]. While the mature dry beans of P. vulgaris and P. lunatus are primarily consumed, P. coccinus is consumed primarily as immature pods (Kay, 1979). Perhaps the most promising aspect of this genus is the fact that there are already well-established breeding programs working to improve bean cultivation. The International Centre for Tropical Agriculture (CIAT) is a leader in Phaseolus breeding, and makes available improved seed accessions, including climbing varieties (Broughton *et al*., 2003).

### Pisum

Pisum sativum, commonly known as green/garden pea, is a temperate or cool season annual climbing plant that has been used by humans since the Bronze Age (Cousin, 1997; Kay, 1979). It likely originated in Ethiopia and Afghanistan before moving to the Mediterranean region and beyond (Cousin, 1997). Branches, frames and nets are all used as climbing supports for its tendrils. There are both spring and winter types of this crop, as well as a wide range of morphologies (Cousin, 1997; PFAF, 2012), made famous by Mendel as the foundation for genetics. The species is generally separated into four subcategories based on end-use: field peas which are used as a forage, market peas used as fresh vegetables for human consumption, vining peas for processing such as freezing and canning, and dried peas which contribute to both human food and animal feed (Cousin, 1997). Similar to P. vulgaris, P. sativum is consumed widely and is an important source of dietary protein in many developing regions (Santalla *et al*., 2011) with a composition high in starch, but also containing between 23-33% protein (Cousin, 1997). P. sativum can be consumed as immature pods, mature peas, sprouts or further processed into secondary products like flour (PFAF, 2012). Newer genetic research is focusing on improving yields by limiting vegetative growth (leaf area), including by converting some leaf growth to tendrils, therefore potentially increasing the climbing strength of the species (Cousin, 1997; Santalla *et al*., 2001). Breeding offers potential to counteract this crop’s susceptibility to pathogens (eg. Fusarium, pea mosaic virus) and drought (which arrests nitrogen fixation) (Cousin, 1997) that terrace farmers may not have the resources to mitigate with expensive inputs.

### Psophocarpus

Psophocarpus tetragonolobis is a twining climbing perennial that is currently only cultivated on a small scale, but shows considerable promise based on its agronomic and nutritional traits (NRC, 1981; PFAF, 2012). It is commonly called winged bean and currently cultivated in humid, tropical environments in Asia and some parts of Africa, though its origin is unconfirmed (NRC, 1981). Typically this species experiences success with 700-4000 mm rainfall annually, however some anecdotal drought resistance has been reported, such as reports from India, as well as from Thailand where this crop survived a severe drought in 1979 while most other crops failed (NRC, 1981). It can grow on a wide range of soils, including those with limited organic matter, relevant for leached terraces, and it is grown almost exclusively by subsistence farmers (NRC, 1981). Winged bean is a valuable green manure due to its exceptional nodulation ability (PFAF, 2012) and has also been successfully grown as a cover crop with tree species such as coconut, banana, oil palm, rubber and cacao in Ghana (NRC, 1981). The crop uses these trees as support to climb without inhibiting their growth, and otherwise requires stakes for support (NRC, 1981). Arguably the most valuable trait of this species is that most organs are edible, including the immature pods (which reportedly can be harvested in as little as 20 days), leaves (rich in vitamin A), shoots (asparagus-like), flowers (used as a garnish or similar to mushrooms), tubers (in certain varieties) and seeds (NRC, 1981; PFAF, 2012). The seeds have almost an identical nutritive value to soybeans, containing a significant amount of protein (around 37.3%), with the advantage of superior palatability (Cerny *et al*., 1971). This feature eliminates the need for fermentation that is required to produce many soy products traditionally consumed, particularly in Asia (Cerny *et al*., 1971). Winged bean flowers under short days, which limits its ability to be cultivated in temperate summers (NRC, 1981), however research is being conducted to develop day neutral varieties (PFAF, 2012). Unfortunately, there is almost no contemporary literature pertaining to agronomic practices associated with this crop. Reviews from the 1970’s (Cerny *et al*., 1971; NRC, 1981) hailed winged bean as being extremely promising, however its continued underutilization indicates that more research is needed to make its adoption a success, especially on terraces.

### Pueraria

Members of the Pueraria genus are commonly referred to as kudzu and are generally aggressively climbing perennials (Keung, 2002; Mitich, 2000). Three main species are of interest when considering agricultural endeavours: P. montana (common kudzu), P. phaseoloides (tropical kudzu), and P. tuberosa (Indian kudzu). These species are distributed throughout Asia and Oceania (Keung, 2002). P. montana was introduced to the United States in 1876 for use as a cover crop, whereas P. phaseoloides was spread throughout tropical regions of Africa, Asia and America for the same purpose (Keung, 2002). The climbing trait of these species allow them to grow upwards on supports or spread along the ground (Keung, 2002). There is limited agronomic data available for this genus, and most of what is reported pertains to the species P. montana. The Pueraria species appear to be adapted to many adverse conditions, such as drought, acidic and marginal soils, and recently disturbed or depleted land, however they cannot tolerate waterlogging (Keung, 2002; Mikhailova *et al*., 2013; Tsugawa, 1985). It has been reported that P. phaseoloides is suitable for growth as a cover crop under coconut, showing exceptional nodulation and nitrogen fixation, and shows potential for intercropping with other plantation crops (Keung, 2002; Thomas and Shantaram, 1984). P. montana and P. phaseoloides have both been noted to be used as a forage and cover crop, and P. tuberosa roots can be eaten raw as a famine food or used as an animal feed, but it is also reported to have a multitude of traditional medicinal uses (Keung, 2002; PFAF, 2012). P. montana can also be consumed by humans, either the cooked roots or young shoots and leaves (PFAF, 2012). It has been reported that kudzu should not be harvested or grazed in the first two years of growth to prevent failure, however once established it produces well for grazing and can recover from livestock trampling and defoliation (Tsugawa, 1985). In fact, kudzu can become so competitive that the crowding out of other crops may become an issue, and its cultivation has been discouraged in the United States since 1950 due to damage caused by its climbing of buildings and telephone lines (Keung, 2002; Mitich, 2000). This vigorous growth can also makes mechanical cutting or mowing of kudzu difficult (Tsugawa, 1985). These issues seem to have limited the further exploration of Pueraria species for agriculture, and recent studies concerning its agronomic details are rare. The crop may be ideal for planting along the base of especially tall terrace risers.

### Sphenostylis

Though there are seven species of this genus that grow in the dry forests and savannas of Africa, by far the most economically important, widely distributed and morphologically diverse species is Sphenostylis stenocarpa. A hallmark of this African species is that produces both edible grains and tubers. As a result, this legume is informally referred to as African yam bean, but it is a little-known crop that nevertheless holds importance to tropical Africa (NAS, 1979). It is cultivated deliberately throughout much of western Africa, but is gathered from the wild in other parts of the continent, with most production being based on traditional indigenous knowledge (Oagile *et al*., 2007; Potter, 1992). When grown deliberately, it is often harvested as an annual, however if its tubers are left in the ground they can act as organs of perennation (tuber-based regrowth) (Potter, 1992). It has been reported that it requires a humid tropical environment with well-drained soils to be successful, but can tolerate acidity and low-fertility reasonably well (NAS, 1979). It is normally grown on trellises or stakes, and varieties vary in their climbing ability from delicate to robust (Oagile *et al*., 2007; NAS, 1979). Some reports claim that one may still yield tubers from unsupported S. stenocarpa plants (NAS, 1979). This crop has also been noted to have a high capacity to deposit nitrogen for subsequent crops, for it has a low N harvest index, which shows potential for its use as a green manure or cover crop (Oagile *et al*., 2007).

As noted above, the main use of this species is for human consumption of both the seeds and tubers (Potter, 1992; NAS, 1979). The tubers take 7-8 months to mature and are reported to have a flavour similar to potatoes, however a much higher protein content (Potter, 1992; NAS, 1979). The seeds mature in a similar timeframe, and must be soaked and/or boiled for several hours to soften, which is often pointed out as a limitation to the crop’s adoption (Potter, 1992; NAS, 1979). The seeds may then be boiled, fried or made into a paste and are reported to contain between 19.5-29% protein (Potter, 1992; NAS, 1979). There is limited mention of the use of S. stenocarpa as animal feed or forage, although one study did note it as a potential good source of protein for livestock (Potter, 1992). Yields have been claimed to be as high as ~2000 kg/ha, but more typically are reported to be around 300-500 kg/ha (Potter, 1992; NAS, 1979). These low yields are the result of several production constraints, which likely also limit adoption of this crop beyond its current range. These constraints include inadequate agronomical guidelines, lack of uniform planting material (for either seed or tuber propagation), and a lack of improved varieties (Oagile etal, 2007).

### Vicia

The genus Vicia contains ~210 species, and has been investigated by The International Center for Agricultural Research in the Dry Areas (ICARDA) for its potential to provide much needed forage to the growing livestock population in West Asia and North Africa (Abd El Moneim, 1993; Raveendar *et al*., 2017). Vicia sativa, or common vetch, is an annual climber that has been noted to grow in this area and has been reported to show good seed yields, herbage and digestibility for use as a forage, though there is considerable variability between varieties (Abd El Moneim, 1993; PFAF, 2012). It has been reported that the early developing fibrous root system and early nodulation make this species suitable for low-input systems, since these nodules can supply nitrogen from the early stages of plant growth (Vlachostergios *et al*., 2011). The seeds of V. sativa are also noted to be nutritious for humans (though not very palatable), which may be cooked or dried and ground into flour (PFAF, 2012). A technique that has been investigated for organic or low-input farming involves planting a mix of several cultivars of V. sativa with the goal of more stable yields and disease resistance; this approach shows promise despite some practical difficulties (Vlachostergios *et al*., 2011). The work conducted on vetches by ICARDA may support the potential cultivation on terrace walls during the dry season.

Another species of some note is V. americana, a perennial sprawling or twining legume with extremely variable morphology that grows throughout North America, from the Yukon and Northwest Territories to Texas and California (Kenicer and Norton, 2008). The cooked young shoots as well as immature pods and mature seeds have been reported to be consumed by indigenous populations of North America (PFAF, 2012). However, for this species to be useful for terrace farmers, significant research would need to be conducted concerning its ability grow in new environments and its utility (PFAF, 2012). Similarly, the plethora of species within Vicia likely hold potential as crops for terrace walls, but more research will be required.

### Vigna

The genus Vigna includes several drought tolerant legumes of critical importance to human societies, and includes climbing varieties of cowpea (V. unguiculata) and rice bean (V. umbellata). Cowpea is an annual crop native to Africa that is considered to have great potential to improve the nutritional status of millions of malnourished people (NRC, 2006). It is grown in areas of Africa, Asia and the Americas and is extremely tolerant to heat and drought, with some cultivars producing grain with less than 300 mm of rainfall (NRC, 2006; Ehlers and Hall, 1997). It is generally cultivated at low altitudes and replaced by common bean higher up, however cowpea is reported to be grown at high altitudes in Kenya and Cameroon (Ehlers and Hall, 1997) indicating that growing cowpea is possible in mountainous regions where the majority of terrace farmers are located. Cowpea has also been reported to produce well in shaded environments, further confirming its suitability for terrace agriculture (Bazill, 1987). Cowpea is often intercropped with sorghum, millets, maize, cassava and cotton, though intensive monocrop systems are present in some places (NRC, 2006; Ehlers and Hall, 1997). Edible parts include the fresh green leaves (which are a good source of iron), green pods and beans, but most commonly the dry grain (NRC, 2006; Sprent *et al*., 2010). An attractive feature of this crop as a food source is that it can be cooked quickly, which is useful in places where there may be cooking fuel shortages (Ehlers and Hall, 1997). Cowpea is also used as a forage, particularly when other species have failed due to drought (NRC, 2006). As a green manure it has been reported to fix up to 70 kg of nitrogen per hectare annually (NRC, 2006), however it has also been noted that in some parts of Africa there can be great variability in nodulation which may limit N fixation (Sprent *et al*., 2010). Though high yields can be achieved in intensive systems, typical subsistence farm yields of cowpea remain low (~100-300 kg/ha) (NRC, 2006). Insect damage, both in the field and in storage, is reported to be the greatest constraint limiting the success of cowpea, and development of resistant varieties is one of the main objectives of genetic improvement for the species (NRC, 2006; Ehlers and Hall, 1997). The research focus on cowpea as well as its impressive stress tolerance suggests that this crop has significant potential for farming on terrace walls.

Rice bean is native to southeast Asia (Andersen, 2012; Saikia *et al*., 1999), and has a twining growth habit allowing it to climb up trees or other crops (Andersen, 2012). It is particularly relevant to this review, since as noted earlier, it is the only legume species that is documented to be cultivated on terrace risers already. Short varieties are reported to be grown on terrace walls to provide food and fodder as well as to protect against erosion in Nepal and India (Andersen, 2012). This species is tolerant of many adverse conditions; it can be established on depleted soils, is reported to remain dormant during drought and then flourish when rains return, and is tolerant to many pests and diseases (Andersen, 2012; Haq, 2011). As a food, the green pods may eaten as a vegetable, but it is more common for the dry seeds to be used as a pulse and cooked into ‘dal’, similar to more common crops such as lentils and pigeon pea (Andersen, 2012; Haq, 2011). The crop residues are a nutritious fodder that are known to increase milk production in cattle (Andersen, 2012). Though not particularly high in crude protein compared to some legumes, the bioavailability of rice bean protein is superior, and there are also significant levels of important nutrients like calcium, iron, zinc and fibre (Andersen, 2012; Saikia *et al*., 1999). This species is also commonly used as a green manure, often intercropped with maize and millets to provide additional nitrogen to the soil and suppress weeds (Andersen, 2012; Khadka and Khanal, 2013). Varieties vary tremendously in the time to maturity, ranging from 60 to 130 days (Haq, 2011). In some regions of Asia, the duration of rice bean allows it to be planted in rotation with paddy rice, improving the soil fertility (Haq, 2011), which is noteworthy as paddy rice is grown on a significant number of terraces worldwide. Rice bean has been the focus of an initiative called Food Security through Ricebean Research in India and Nepal (FOSRIN), and there are large repositories of germplasm in both India and Taiwan, to enable continued development of this crop (Andersen, 2012; Haq, 2011). However, there are some constraints to farmer adoption of rice bean, particularly issues of indeterminate versus determinate growth varieties, and inconsistent seed size and hardness (Andersen, 2012). Continued investment in optimizing agronomic practices and improving varieties would allow terrace farmers to take full advantage of this promising legume.

## 7. Challenges, recommendations and conclusions

### 7.1 Summary of opportunities

The purpose of this paper was to review climbing legumes that have potential to grow on terrace walls to address the challenges faced by millions of subsistence farmers in mountainous regions, including: poor access to inputs (fertilizers, herbicides and pesticides), female drudgery (e.g. the need to climb up and down steep gradients to weed), exacerbated human and livestock malnutrition, and vulnerability to climate change. Though all the crops reviewed show promise, some crops are worth highlighting:

- With respect to input replacement, African yam bean and jack bean have a particularly high capacity to provide nitrogen compared to other crops (Oagile *et al*., 2007; Wortmann *et al*., 2000).
- Jack and velvet bean have been shown to discourage nematodes (Arim *et al*., 2006; Witcombe et al., 2008).
- In terms of drudgery reduction, horse gram has been shown to decrease the labour required for weeding by providing ground cover (Arim *et al*., 2006; Witcombe *et al*., 2008).
- Nutritionally, horse gram, cowpea and rice bean can be significant sources of iron, while rice bean also contains high levels of zinc, and the leaves of winged bean are a good source of vitamin A (Andersen, 2012; Bravo *et al*., 1999; Ehlers and Hall, 1997; Saikia *et al*., 1999; NRC, 1981; Sudha *et al*., 1995).
- In terms of climate change resiliency, cowpea, common bean, kudzu, and green pea are all especially drought tolerant (NRC, 2006; Kay, 1979; Mikhailova et al., 2013).

### 7.2 Potential challenges

With so many potential advantages, the question must be asked as to why most terrace farmers around the world have not adopted the practice of using terrace walls for growing plants, including legumes. The exception, as noted earlier, appears to be rice bean (Andersen, 2012). There may be several reasons for this gap. In particular, the terrace agro-ecosystem itself may present challenges. For example, paddy rice production involves seasonal flooding and would lead to waterlogged soils, which may be incompatible with some climbing legumes. However, three of the reviewed species in particular are reported to perform even in waterlogged soils: A. americana (groundnut), C. ensiformis (jack bean), and L. sativus (grass pea) (Duke, 1983; Haq, 2011; Malek *et al*., 2000). Terrace walls have diverse heights and angles, making cultivation of any one crop variety challenging. If climbing legumes are grown directly on the terrace walls, there may also be pest problems with soil borne pathogens or insects emerging from riser soils. Climbing varieties may intercept sunlight, to shade crops adjacent growing on the horizontal surface of the terrace. Aggressive perennials such as kudzu (which is known to be smothering) would be of most concern (Keung, 2002; Mitich, 2000). Access to the seeds of appropriate varieties, and breeding of climbing varieties for local conditions, represent additional practical challenges. Finally, farmer adoption is a challenge with any innovation, and there need to be clear economic benefits with little additional labour for farmers to be attracted to a new practice. This is particularly true when additional challenges present themselves as in the case of underutilized (and hence under-developed) crops, as evidenced by the dis-adoption of rice bean in the Himalayan region (Andersen, 2012).

### 7.3 Recommendations for the future

We have nine specific recommendations to help accelerate research into farming on terrace walls (FTW), related to overcoming agronomic, breeding and socio-economic challenges:

#### 7.3.1. Agronomy

Local agronomists expertise in terrace agriculture must be recruited to undertake:

1. Field trials on research plots: Using controlled research plots, good quality agronomic data will be required for the climbing legumes such as optimal seeding rates and spacing, as well as to understand interactions (competition and synergies) with the crops already grown on the horizontal terrace surface. Trials will need to be conducted to find species most suitable for local climactic and soil conditions, and fit them appropriately into local cropping calendars.
2. Establishment of best practices for farmers: Once agronomic trial data has been collected, that information must be transferred into practical instructions for farmers. For farmers to adopt a new crop variety and practice, they will need to know how, when and what to cultivate. Management practices for pests as well as any fertilizer recommendations will be required, with a deliberate focus on low chemical inputs. Legumes require compatible rhizobia to be present in the soil to fix nitrogen, which may limit farmer adoption of a new legume species, or require introduction of appropriate microbial inoculants. If tendril strength or the riser material is inadequate or unsuitable, the introduced climbing legumes may require trellises made from local wood resources. Finally, to ensure that nutrients and water are sufficient at the base of the terrace risers, the terraces may need to be inverse-sloped.
3. On farm systems-level evaluation: Using on-farm split plots and participatory approaches, the real world benefits and challenges of FTW must be evaluated side by side with conventional terrace farming at a systems-level, focusing on agronomic changes (changes in yield, soil quality) and socioeconomic indicators (nutrition, income, labour, resiliency).

#### 7.3.2. Breeding

Participatory plant breeding may help ensure that the specific needs of the local agricultural system are addressed to ensure farmer adoption of improved varieties. We have three recommendations in this area:

1. Breeding to suit the terrace microenvironment: Candidate climbing legumes may need to bred for improved climbing ability, improved shade resistance, tolerance to waterlogging and shorter duration.
2. Breeding for a low-input system: To adapt climbing varieties to a low input system, characteristic of terrace farms, breeding will be required for improved N fixation, pest resistance, and tolerance to abiotic stresses.
3. Overcoming crop specific limitations: Some of the species suggested in this review have particular traits that require breeding to improve their use on terrace walls. For example, P. montana (kudzu) requires breeding to make it less aggressive, while Lathyrus species require breeding to decrease neurotoxin levels.

#### 7.3.3. Socio-economic factors

1. Local availability, acceptability and rates of adoption: A critical step towards the success of FTW would be ensuring that seeds of climbing varieties would be available to remote terrace farmers, with no regulatory or physical restrictions. Co-operation between governments, private seed companies and centres of the Collaborative Group of International Agricultural Research (CGIAR) would help to overcome barriers such as varietal approval. For seed distribution into remote regions, using existing networks such as snack food and alcohol vendors may be useful. Participatory approaches will ensure local acceptability of taste, texture, look and quality of new food products. Long-term evaluations will be needed to track farmer adoption, successes and challenges of FTW.
2. Extension: It will be challenging to disseminate the FTW concept and provide training in remote areas at a reasonable cost. The SAK Nepal project (Chapagain and Raizada, 2017) has begun experimenting with captioned picture books to demonstrate the technique (Figure 3), and these resources are open access and can be downloaded for free.
3. Development of markets: To provide market incentives, value chains will need to be established to permit sales of surplus crops for human food or animal feed and forage.

**Figure 3.**
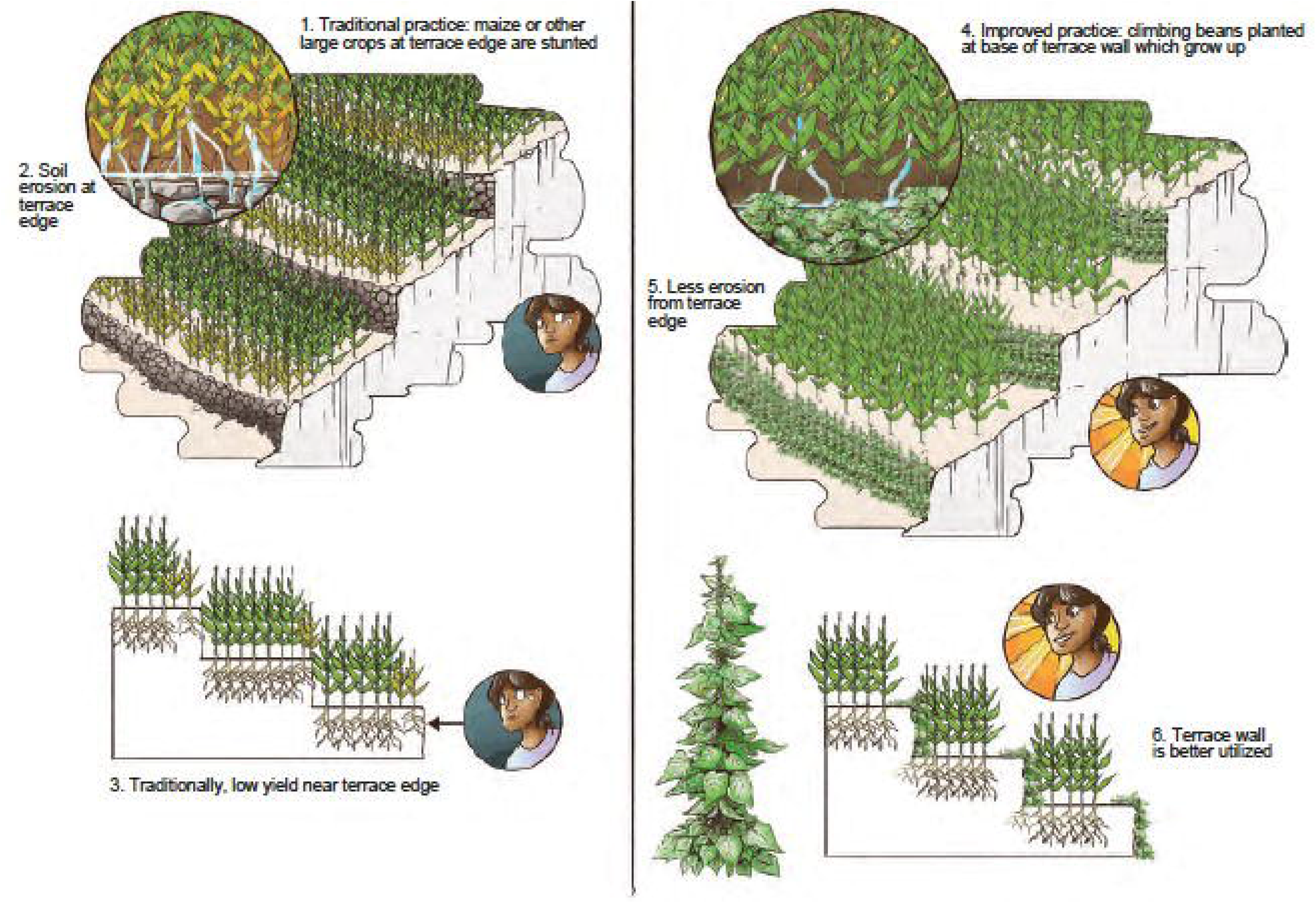
Agriculture extension lesson to train smallholder farmers about the potential of growing climbing legumes on terrace risers. Image courtesy of Lisa Smith, University of Guelph, can be can be reused under the Creative Commons BY licence

### 7.4 Conclusions

It is hoped that this review has helped to shed light on the specific challenges faced by terrace farmers around the world, and provided an avenue for future innovation in these ancient farming systems. There are many candidate climbing legumes that have potential to grow on terrace walls. Many of these crops are nutritious to humans and/or livestock, can grow under low input conditions and show tolerance to drought and shade. This review has noted the practical challenges that may limit farmer adoption of FTW, but we hope that these may be addressed by the concrete recommendations listed. In an era where human population growth will occur primarily in developing nations at a time of unpredictable climate change and environmental degradation, growing legumes on terrace walls provides an opportunity to reduce the increasing vulnerabilities of terrace farmers.

## Acknowledgements

This research was supported by a grant to MNR from the International Development Research Centre (IDRC) and Global Affairs Canada as part of the Canadian International Food Security Research Fund (CIFSRF).

## Author contributions

MNR and JCC conceived the manuscript. JCC wrote the manuscript and MNR edited the manuscript.

